# Informeasure: an R/Bioconductor package to quantify nonlinear dependence between variables in biological networks from an information theory perspective

**DOI:** 10.1101/2021.12.20.473524

**Authors:** Chu Pan, Limei Jing, Jiawei Luo, Xiangxiang Zeng

**Affiliations:** College of Computer Science and Electronic Engineering, Hunan University, Changsha, Hunan, China

**Keywords:** R/Bioconductor package, information measure, nonlinear dependence, biological regulatory network

## Abstract

Using information measures to infer biological regulatory networks can observe nonlinear relationship between variables, but it is computationally challenging and there is currently no convenient tool available. We here describe an information theory R package named Informeasure that devotes to quantifying nonlinear dependence between variables in biological regulatory networks from an information theory perspective. This package compiles most of the information measures currently available: mutual information, conditional mutual information, interaction information, partial information decomposition and part mutual information. The first estimator is used to infer bivariate networks while the last four estimators are dedicated to analysis of trivariate networks. The base installation of this turn-key package allows users to approach these information measures out of the box. Informeasure is implemented in R program and is available as an R/Bioconductor package at https://bioconductor.org/packages/Informeasure.

## Introduction

Quantifying dependence within variables in biological networks is an important issue since understanding the condition-responsive activations of biological process hinges on the accurate construction of biological network[1]. There are several computing criteria that are used to quantify regulatory relationship between variables in biological regulatory networks[2]. Correlation methods such as Pearson correlation coefficient and partial correlation are widely employed to assess the linear relationship between paired variables. Information measures as alternatives can be used to evaluate the dependence between two variables or even multiple variables with considerable advantages over correlation methods, as they are able to capture more general nonlinear associations and reflect dynamics between variables. Most information measures currently available includes mutual information (MI), conditional mutual information (CMI)[3], interaction information (II)[4], partial information decomposition (PID)[5] and part mutual information (PMI)[6]. All these estimators have applications in bioinformatics, particularly in the inference of triplet regulatory networks from expression profile data. Chen and colleagues employed CMI algorithm to reconstruct gene regulatory network, and on this basis developed a new multivariate information theory measure, known as PMI algorithm[6-8]. Califano and colleagues used multivariate information measures like CMI and II to simulate the regulatory process of different types of RNAs competing for the same miRNAs, and to evaluate the regulatory effect of RNAs in modulating the other miRNAs by calculating the changes of their abundance, hence the identification of the competing endogenous RNAs regulatory networks[9-11]. Stumpf and colleagues applied the PID to single-cell data to evaluate the statistical relationships within triplet variables in gene regulatory network[12, 13]. Although information measures such as PMI provided source code in the paper, as did network computational methods like Hermes[9] and PIDC[12], they were implemented in different programming, which limits the widespread promotion of these algorithms in bioinformatics. R has become one of the most widely used programs in bioinformatics due to its data handling and visualizing abilities as well as its flexibility. There are currently some R-code toolkits for estimating information metrics such as R package minet developed by Meyer *et al*.[14] and entropy written by Hausser and Strimmer[15], both aimed at solving the problem of inferring gene association networks from high-dimensional expression profile data. But their most powerful functions can only infer two-variable regulatory relations by mutual information. In order to leverage information measures on more complicated regulatory network inferences, we expand entropy to a more comprehensive tool called Informeasure. This R package integrates five information measures as mentioned above and provides easy-to-reach interfaces for users. The base installation of Informeasure allows users to approach most information measures directly, and offers a basis for creating novel methods for situations not covered by existing methods.

## Information theoretic measures

Before introducing the proposed R package, we briefly describe the principles of these five information measures so that based on the description users can choose an appropriate measure for their specific purposes. The first one is MI, used to measure the mutual dependence between two joint random variables. It evaluates the expected amount of information obtained about a variable by observing the other variable. The notion of mutual information is derived from the entropy of random variables. While the entropy is a fundamental concept in information theory that quantifies the amount of uncertainty contained in a random variable. In the discrete case, the formula for entropy is given by:

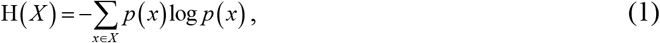

therefore, the MI value of two jointly discrete variables *X* and *Y* can be computed as a double sum of probabilities, or equivalently expressed as a chain of entropies:

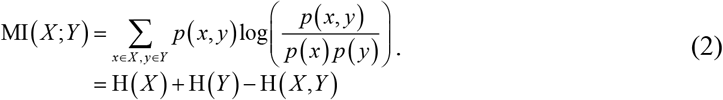

If the third variable *Z* is imposed on the two joint discrete variables *X* and *Y*, the mutual information is conditioned upon a third variable to yield the concept of CMI. The information measure CMI is widely used to evaluate the expected mutual information between two random variables conditioned on the third one. For the discrete variables *X, Y* and *Z*, the formula for CMI is as follows:

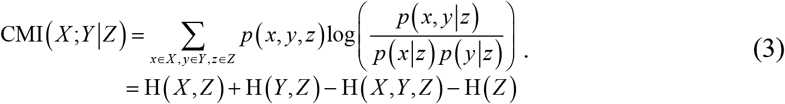

PMI is another recently developed information criterion that is devoted to measuring the nonlinear direct dependence between two random variables given a third, especially if any one variable has a potentially strong correlation with the third one. Again for the discrete variables *X, Y* and *Z*, the formula for PMI is given by:

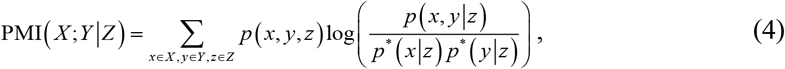

where *p*(*x, y* | *z*) *p*^*^(*x* | *z*) *p*^*^(*y* | *z*) is the partial independence of discrete variable *X* and *Y* given *Z*. Here *p*^*^(*x* | *z*) and *p*^*^(*y* | *z*) are defined as:

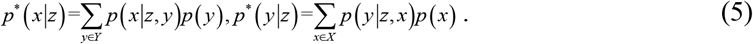

Another type of multivariate information measure is II, also known as co-information, which measures the amount information (synergy or redundancy) contained in a set of variables beyond any subset of those variables. The II value can be either negative or positive. Here in the case of three variables, a negative II value indicates that the third variable inhibits the correlation between the first two variables, while a positive II value indicates that the third variable promotes or strengthens the correlation between the first two variables. The formula for II can be simply expressed as:

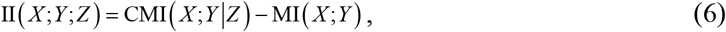

PID is a new emerging branch of information theory, which decomposes source information acting on the target into four parts: joint information (synergy), individual information (two unique), and shared information (redundancy). Here again in the case of three discrete variables *X, Y* and *Z*, the formula for PID is as follows:

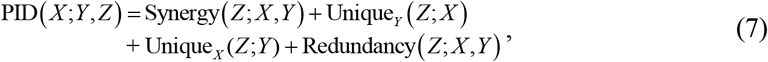

The synergy is the bonus information about *Z* provided by variables *X* and *Y* together not by individual variable, while the redundancy refers to the portion of information about *Z* that provided by either variable *X* or *Y* alone. The unique contribution from *X* (or *Y*) is the part of information provided only by *X* (or *Y*). Here first considers the specific information provided by *X* about the target variable *Z* in state *z* :

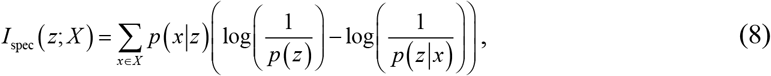

Let *S* ={*X, Y*} and a target variable *Z*, the redundant information is evaluated by comparing the amount of information provided by each variable within *S* about each state of the target variable *Z*,

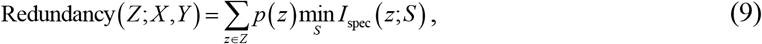

The unique information terms can be calculated based on the relationship between redundant information and the pairwise mutual information, as shown below:

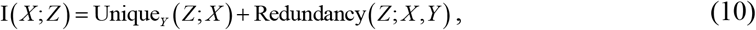

Finally, the synergistic information is evaluated through the interaction information, because the interaction information is the difference between the synergistic and redundant information:

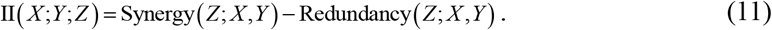

In summary, MI that is applicable to quantify nonlinear dependence between two variables in biological networks, while the characteristics of CMI and PMI are fully suitable for trivariate network inference. The synergistic and redundant information decomposed from II and PID can help biologists better decipher the cooperative and competitive relationships that occur in biological networks.

### Implementation and main functions

Information measure is typically implemented by first discretizing continuous variables into a count table, evaluating probability from the counting, and/or then estimating entropy according to the (joint) probability matrix, finally calculating the information value that is the most representative for the association between variables. As such the package breaks the entire implementation into three main parts: data discretization, probability/entropy estimation and information estimation. This user-oriented implementation allows users to freely combine discretization methods, probability or entropy evaluators and information metrics according to their specific requirements and purposes. In detail, two of the most common discretization methods are adopted in this package. One is a uniform width-based method (default), which divides the continuous data into *N* bins with equal width. The other alternative is a uniform frequency-based approach, which determines the continuous data into *N* bins with equal count number. By default in both methods, the number of bins is initialized into a round-off value according to the square root of the data size: 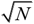. In the process of probability estimation, three types of probability estimators referencing to the entropy package are available named the empirical estimator (default), the Dirichlet distribution estimator and the shrinkage estimator, while the Dirichlet distribution estimator also includes four different distributions with different prior values. They are listed as follows:

✧ method = “ML”: maximum likelihood estimator, also referred to empirical probability;
✧ method = “Jeffreys”: Dirichlet distribution estimator with prior a = 0.5;
✧ method = “Laplace”: Dirichlet distribution estimator with prior a = 1;
✧ method = “SG”: Dirichlet destruction estimator with prior a = 1/length(*XY*), where *XY* is the joint count table for variables *X* and *Y*;
✧ method = “minimax”: Dirichlet distribution estimator with prior a = sqrt(sum(*XY*))/length (*XY*);
✧ method = “shrink”: shrinkage estimator.

However, the most important functions in this package are five different information measures, all of which end with .measure() in form. They are MI.measure() for MI, CMI.measure() for CMI, II.measure() for II, PID.measure() for PID and PMI.measure() for PMI. In the simplest form, each function can be called with only a joint count table. But each function also provides six available probability estimation methods as listed above and three different base logarithmic calculations for users to choose. All functions except PID.measure() return a numeric value that represents the information measurement between two variables or among three variables. The PID.measure() function returns a list that includes synergistic information, unique information from one source variable, unique information from the other source variable, redundant information, and the sum of the four parts of information.

### Application in transcriptome regulatory network inferences

We consider estimating information measures from breast cancer expression profile data generated by The Cancer Genome Atlas (TCGA), which is the sample data stored in the ‘extdata’ directory, with applications in various types of transcriptome regulatory network inferences, as shown in Figure. 1. In the case of two variables, we apply MI to identify the dependence between proteins in protein-protein interaction network inference. Figure. 1A shows the MI value of BRCA1 and BARD1 is the highest among all pairs, which agrees with the evidence that these two proteins come from a protein complex and are with high dependence on each other. In the three-variable case, we apply CMI and PMI to the triplet network inference, as both measures claim to be able to evaluate the influence of the third variable on the mutual information of the two joint variables. Such characteristics of these two information measures are fully applicable to the ceRNA network inference. Taking lncRNA-associated competitive triplet as an example, the calculation is the perturbation intensity on miRNA-mRNA by lncRNA. As can be seen from Figure. 1B, the result value of PMI is higher than that of CMI. This is because PMI can accurately evaluate the dependence between hsa-miR-26a-5p and PTEN even though PTENP1 has a strong correlation with PTEN as is illustrated in the linear fits in Figure. 1B, whereas CMI tends to underestimate the result in this case. Here again in the three-variable case, we apply II and PID to explore the cooperative or competitive regulation mechanism of two miRNAs on the common target mRNA, given that they can extract synergetic or/and redundant information between variables. As shown in Figure. 1C and 1D, the positive value of II and the high synergy value of PID mean that the two miRNAs together can provide most information for the target variable, implying the cooperation mechanism of the two miRNAs. The boxplot shows that the expression patterns of hsa-miR-34a-5p and hsa-miR-34b-5p are similar but different from the target MYC, which is consistent with the fact that the two miRNAs coordinately regulate the common target mRNA

**Fig. 1.**
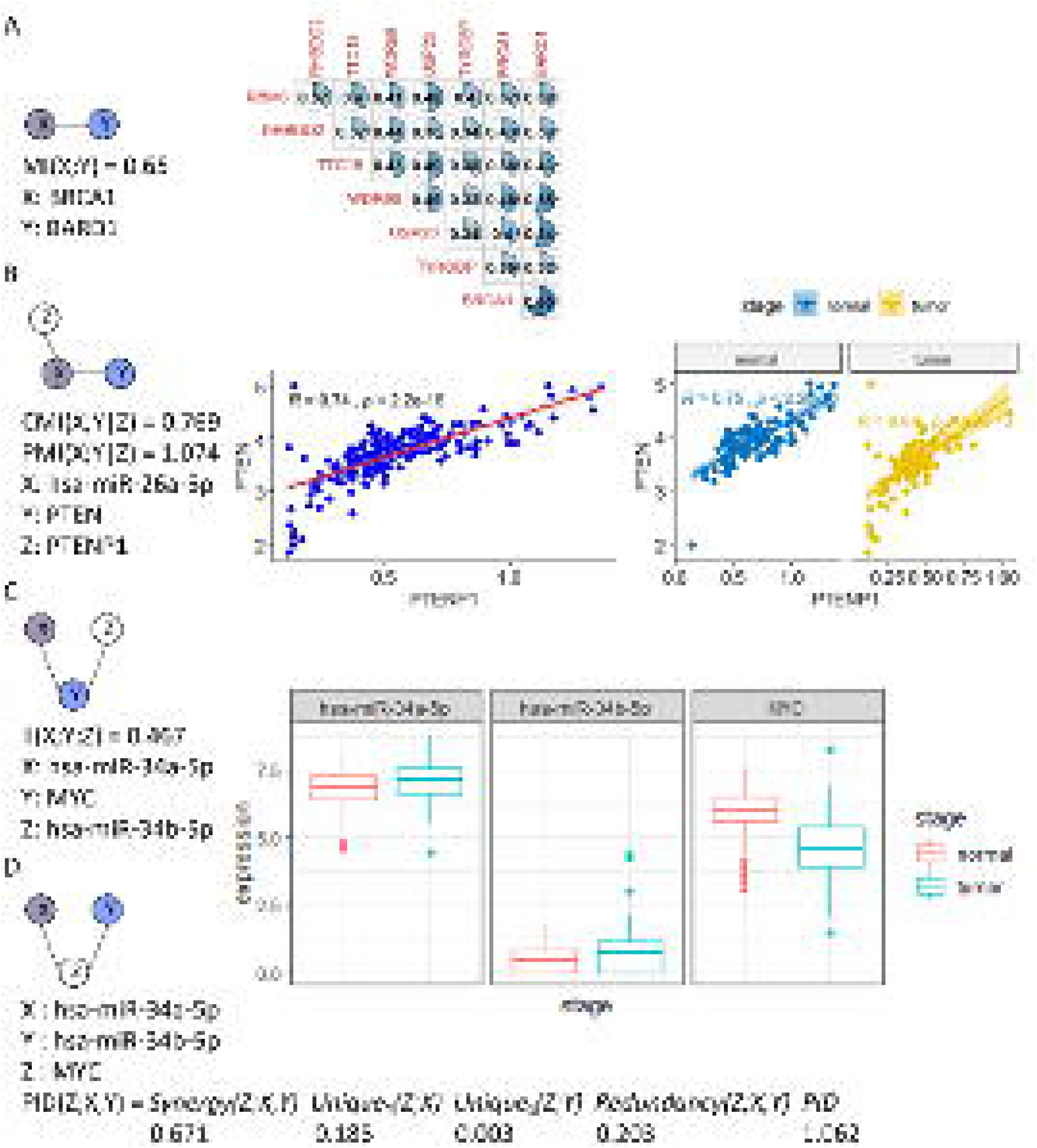
Application scenarios of five different information measures. **(A)** Mutual information is used to assess the association between proteins. Here RNA-Seq data are used to approximate protein expression data. **(B)** Conditional mutual information and part mutual information are employed to identify lncRNA-associated ceRNA network. **(C)** Interaction information is utilized to determine miRNA cooperative/competitive regulatory network. **(D)** Partial information decomposition is utilized to evaluate the miRNA cooperative/competitive regulatory network as well. The results obtained were calculated according to the default parameters. Graphs illustrated above were created by R packages corrplot, ggplot2 and patchwork.

### Test on large-scale data

In this R package, we are responsible for providing users with easily accessible interfaces to approach five information measures. Users can read the walkthrough examples described in the vignette as a reference. In general, much small data runs fast and well on a single processor, but thing will be bad once the amount of data reaches a certain point because of the limitations of the single processor’s computing power. While estimating all possible interactions and dependencies in large-scale networks within a reasonable time is a critical issue. At this time, we suggest that users add an R built-in package named parallel to speed up the operation, as each task of calculating the nonlinear relationship between variables is completely independent of the others, thus parallel execution of these tasks is the preferred option. We again consider using the expression profile data generated by TCGA to test the feasibility of Informeasure being applied to large-scale data. We choose breast carcinoma (BRCA) and thyroid carcinoma (THCA), as these two data are in different sample sizes. The BRCA and THCA data include 1181 and 558 samples, respectively, each measuring the expression of multiple lncRNAs, miRNAs and mRNAs. The expression data are further log2-transformed. The machine on our hardware platform has 2 physical CPUs, each of which includes 8 processing cores with 16 hyper-threads, a total of 32 threads are used for computation. The CUP configuration is as follows: Intel(R) Xeon(R) Silver 4208 CPU @ 2.10GHz. We start with 1 thousand gene pairs/triplets, and gradually increase the data size to 1 million at 10-fold intervals. Gene pairs/triplets (mRNA-mRNA for MI, lncRNA-miRNA-mRNA for CMI and PMI, mRNA-miRNA-mRNA for II and PID) are sampled randomly from the genes (i.e., lncRNA, miRNA and mRNA). We employ the parallel::mclapply function to send tasks encoded as function calls to each processing core on our machine in parallel. We use R built-in function system.time() to test the time consuming, resulting in ‘user’, ‘system’ and ‘elapsed’, where the ‘elapsed’ is the wall clock time spent executing the calculation. As illustrated in Figure. 2, the CPU time consumption increases with the increase in the number of gene pairs/triplets and the sample size of expression profile data, but all tasks are completed within an acceptable time. Among these five information measures, PMI seems to be the most time-consuming, while even on a million BRCA data, it can be done in 1.5 hours. All other tests on 1 million data set are completed within an hour, showing the effectiveness of Informeasure in computing power. Of course, parallelization is greatly beneficial for processing such large-scale data.

**Fig. 2.**
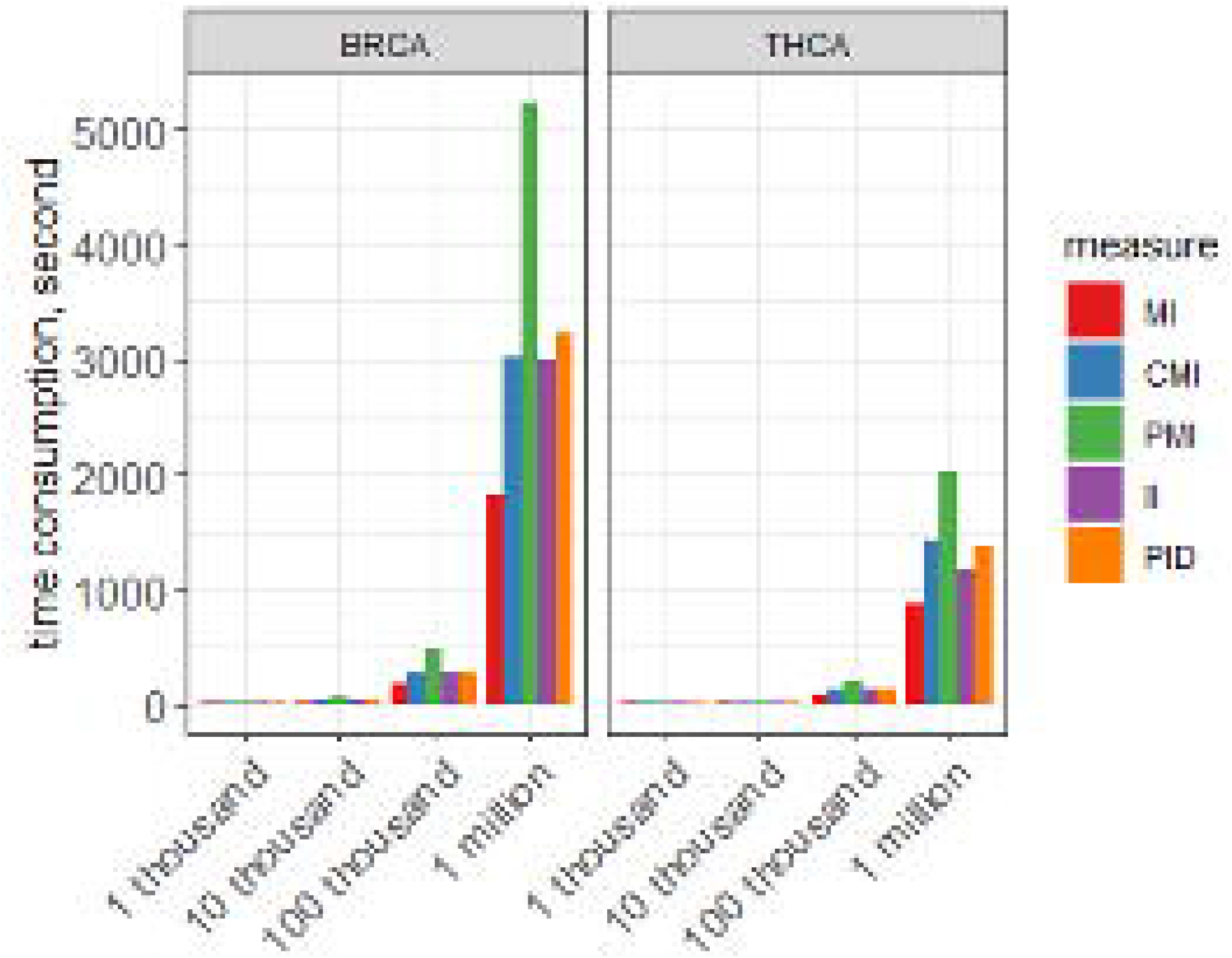
Time consumption results of five different information measures tested on different datasets.

## Discussion and conclusion

Our goal is to provide researchers with a comprehensive tool of information measures to help them address specific research purposes. The developed tool Informeasure is an R/Biocondutor package with documented functions and demonstration examples described in the vignette, so users can easily access these information measures. In order to consider the efficiency of the code, we use vector/matrix calculation as much as possible in the implementation to avoid unnecessary loops. Even when loops are unavoidable, we employ optimized functions such as apply and lapply instead of for and while. In this current version, we are mainly focus on applying information measures to two- and three-variable cases, although measures such as II and PID claim to be extended to higher dimensions. But to the best of our knowledge identifying nonlinear dependence between two- and three-variable is currently the main concern, so we choose three variables as the largest network unit handled by this toolkit. In addition, biologists are often asked to verify the statistical significance of the values obtained from information measures. The classic approach is to first generate a null hypothesis by measuring the distribution of multiple pairs/triplets created under random conditions, and then evaluate the significance of each pair/triplet form the null hypothesis model. Our package does not provide statistical analysis, as the distribution generated under random conditions is uncertain, but the creation of the distribution can be done quickly according to the parallel strategy mentioned above. In conclusion, we provided the implementation of five information measures in this R package. The brief survey of information theory algorithms guided users to choose appropriate measures for specific purposes. The illustration successfully surveyed information measures for inferring various types of regulatory networks from expression profile data, with primary focus on trivariate networks. The computing power test demonstrated the capability of Informeasure being applied to large-scale data. We are convinced that Informeasure can be widely serving as a useful tool to facilitate inferring complex condition-specific regulatory networks. Our goal is to clarify these methods for other researcher as they search for the multivariate information measure that will best address their specific research goals. We are not attempting to guide the development of information theory; rather we seek to promote the wider use of information theoretic tools.

## Acknowledgements

The authors thank Ms. Song Jing for her careful proofreading of the manuscript, Mr. Xianghua Wang for his helpful discussions on the PMI algorithm, and Dr. Junpeng Zhang, Mr. Nitesh Turaga and Mr. Martin Morgan for their informative suggestions on writing the R package. C. Pan would like to thank his family for their persistent support during his most difficult times in 2020.

